# TMEM2 Deficiency Causes Cleft Palate and Disrupts Vascular Development

**DOI:** 10.64898/2026.11.20.768019

**Authors:** Hisham Shamma, Yuichiro Nakanishi, Jun-Ichi Sasaki, Hirohiko Sakai, Tomoya Murotani, Taisei Tomohiro, Hiroshi Kurosaka, Yu Yamaguchi, Takashi Yamashiro, Toshihiro Inubushi

## Abstract

Hyaluronan (HA) is a major component of the extracellular matrix during craniofacial development. Studies of hyaluronan synthase 2 (HAS2) have established that HA synthesis is required for normal palatal development. However, the contribution of HA degradation to this process remains poorly understood. TMEM2 is a cell-surface hyaluronidase, and previous studies have demonstrated its role in neural crest cell (NCC) migration and the survival of post-migratory NCCs. Previous Wnt1-Cre-mediated deletion of Tmem2 caused early embryonic lethality, preventing assessment of later palatal development. Here, we used Sox10-Cre to delete Tmem2 in the NCC lineage and investigate its role in palatogenesis. Conditional knockout (CKO) embryos developed complete cleft palate. Palatal shelves elevated but remained smaller than controls, with reduced proliferation and increased apoptosis. ZsGreen lineage labeling revealed reduced directional organization of NCC-derived cells, accompanied by widespread HA accumulation throughout the palatal mesenchyme. To identify molecular changes accompanying these abnormalities, we performed RNA sequencing, which revealed enrichment of hypoxia-related transcriptional programs and suppression of cell-cycle programs. Increased HIF-1α and pimonidazole labeling provided complementary evidence of tissue hypoxia. Vascular analyses revealed altered CD31-positive vessel architecture, reduced contact between NCC-derived cells and vessels, and fewer PDGFRβ-positive cells. Together, these findings identify TMEM2-dependent HA remodeling as an important component of palatal development and link its disruption to altered NCC-derived mesenchymal organization, vascular and perivascular abnormalities, and tissue hypoxia.

## Introduction

Cleft palate is a common craniofacial birth defect resulting from disruption of the coordinated growth, elevation, and fusion of the bilateral palatal shelves (Bush and Jiang 2012; Dixon et al. 2011). The palatal mesenchyme is largely derived from cranial neural crest cells (NCCs), which contribute extensively to craniofacial tissues(Le Douarin and Kalcheim 1999; Minoux and Rijli 2010). During palatogenesis, the growth and organization of NCC-derived mesenchymal cells must be coordinated to support palatal shelf development.

The extracellular matrix (ECM) provides a dynamic microenvironment that regulates cell adhesion, migration, and differentiation during embryonic development(Rozario and DeSimone 2010). Hyaluronan (HA), a major glycosaminoglycan of the embryonic ECM, contributes to tissue hydration, extracellular space formation, and cell–matrix interactions (Toole 2004). The developmental importance of HA is demonstrated by the severe cardiac and vascular abnormalities and embryonic lethality observed in Has2-null mice (Camenisch et al. 2000). Neural crest-specific deletion of *Has2* also causes craniofacial abnormalities, including cleft palate, establishing a role for HA synthesis in palatal development (Lan et al. 2019). However, compared with HA synthesis, the role of extracellular HA degradation in palatal development remains poorly understood. TMEM2 is a cell-surface hyaluronidase that degrades high-molecular-weight HA under neutral conditions (Yamamoto et al. 2017) and promotes integrin-dependent cell adhesion and migration through local removal of pericellular HA (Irie et al. 2021). These findings suggest that TMEM2-mediated HA remodeling may regulate the microenvironment surrounding NCC-derived cells during palatogenesis, but how this process contributes to palatal development remains unclear.

Previous studies using *Wnt1-Cre*-mediated deletion of *Tmem2* in NCCs demonstrated severe craniofacial and cardiovascular abnormalities accompanied by embryonic lethality around E12.5, precluding analysis of its role during later palatal development (Inubushi et al. 2022). To overcome this limitation, we used a *Sox10-Cre* driver to delete *Tmem*2 in NCC-derived cells and thereby examine TMEM2 function during palatogenesis. We investigated whether TMEM2-dependent HA remodeling contributes to palatal shelf growth and the spatial organization of NCC-derived mesenchyme, and whether disruption of this extracellular environment is associated with abnormalities in vascular organization and tissue oxygenation.

## Materials and methods

### Animals and embryo collection

Animal experiments were approved by the Institutional Animal Care and Use Committee of the Graduate School of Dentistry, The University of Osaka (Approval No.30-008-0). Genetically modified animal experiments were approved by the institutional biosafety committee (04263 and 04315). Conditional *Tmem2* floxed mice (*Tmem2^f/f^*) were generated as previously described (Inubushi et al. 2022). Mice were maintained on a C57BL/6J background. *Tmem2^f/wt^; Sox10-Cre* males were crossed with *Tmem2^f/f^* females to generate *Tmem2^f/f^; Sox10-Cre* embryos (CKO), with *Cre-negative* littermates used as controls unless otherwise specified. ZsGreen reporter mice (JAX #007906) were used for lineage tracing. The day of vaginal plug detection was designated E0.0. Embryonic sex was not determined.

### Histological and morphometric analyses

Gross craniofacial morphology was examined at E16.5 using an SZX16 stereomicroscope (Olympus). For palatal assessment, mandibles were removed, specimens were stained with DAPI (1 μg/mL; Fujifilm Wako) for 10 min and imaged using a BZ-X700 fluorescence microscope (Keyence).

Embryonic heads were fixed in 4% PFA at 4 °C for 24 h, cryoprotected in 10%, 20%, and 30% sucrose, embedded in Tissue-Tek O.C.T. Compound (Sakura Finetek), and coronally sectioned at 12 μm. H&E-stained sections were examined at E13.5, E14.0, and E15.5. Palatal shelf length and cross-sectional area were quantified using ImageJ (NIH) on anatomically matched coronal sections selected based on the position of the molar tooth germs.

### Cell proliferation and apoptosis analyses

Cell proliferation was assessed by Ki-67 immunostaining (1:200; Abcam, ab15580) and quantified as the proportion of Ki-67-positive/DAPI-positive cells. Apoptosis was assessed using the In Situ Cell Death Detection Kit (Roche, 11684795910) and quantified by TUNEL-positive cell counts. Ki-67 and TUNEL analyses were performed at E13.5, E14.0, and E15.5. For each assay, equivalent anatomical regions were analyzed between control and CKO embryos.

### HA distribution and NCC lineage analyses

NCC-derived cells were visualized using *Sox10-Cre; ZsGreen* reporter embryos. E13.5 coronal sections were stained with biotinylated HABP (1:100; EMD Millipore, 385911), followed by streptavidin–rhodamine (1:500; Jackson ImmunoResearch, 016-290-084) and DAPI. Images were acquired using identical microscope settings for all samples within each experiment. HABP-positive area was quantified within the palatal mesenchyme. For spatial analysis, the mesenchymal ROI was divided into equal medial and lateral halves, and the medial-to-lateral ratio of HABP-positive area was calculated. ZsGreen-positive cell organization was assessed by structure tensor analysis after Otsu segmentation, with orientation coherence ranging from 0 (random) to 1 (highly aligned). The same segmentation and analysis parameters were applied to all samples.

### RNA sequencing and gene set enrichment analysis

E14.0 bilateral palatal shelves from each embryo were pooled as one biological sample. Total RNA was extracted using the RNeasy Plus Mini Kit (Qiagen), and samples with an RNA integrity number >8 were used for library preparation. Stranded mRNA libraries were prepared using the TruSeq Stranded mRNA Library Prep Kit (Illumina) and sequenced on a NovaSeq 6000 with 150-bp paired-end reads. Reads were aligned to the mouse reference genome (mm39), and differential expression was analyzed using DESeq2. Differentially expressed genes were defined using an absolute log2 fold-change threshold of >0.5. One CKO sample showing a divergent global gene expression profile was excluded from downstream analysis. GSEA was performed using GSEA 4.4.0 and mouse Hallmark gene sets to compare control and CKO palatal tissues. DESeq2-normalized expression data and phenotype labels were used as input, and genes were ranked within GSEA using the signal-to-noise metric. Statistical significance was assessed using gene-set permutation, and enrichment was evaluated using normalized enrichment scores (NES).

### Tissue hypoxia analyses

Tissue hypoxia was assessed at E13.5 by HIF-1α immunofluorescence (1:200; Cell Signaling Technology, #48085S) and pimonidazole labeling. Pregnant mice received pimonidazole hydrochloride (Hypoxyprobe; 60 mg/kg, intraperitoneally), and embryos were collected 1 h later and stained with RED549-conjugated anti-pimonidazole antibody (1:50). HIF-1α- and pimonidazole-positive cells within the palatal mesenchyme were counted. For each assay, equivalent anatomical regions were analyzed between control and CKO embryos.

### Vascular and perivascular analyses

Vascular and perivascular organization was examined in E13.5 palatal tissues. *Sox10-Cre; ZsGreen* sections were immunostained for CD31 (1:200; BD Biosciences, 550274), and the proportion of ZsGreen-positive cells contacting CD31-positive vessels within the palatal mesenchyme was quantified. A ZsGreen-positive cell was considered vessel-associated when its signal was in direct contact with the CD31-positive vascular boundary. For perivascular analysis, sections were co-immunostained for PDGFRβ (1:200; Cell Signaling Technology, #3169) and CD31 (1:400), and PDGFRβ-positive cells were counted per 30,000 μm² of palatal mesenchyme. For three-dimensional vascular imaging, E13.5 tissues were processed using the MACS Clearing system (Miltenyi Biotec), stained with directly conjugated anti-CD31 Vio R667 antibody (1:200; clone REA784; 130-128-736), and optically cleared before imaging under identical acquisition settings between groups.

### Statistical analysis

Data are presented as mean ± SD or as box-and-whisker plots, as specified in the figure legends. Statistical analyses were performed using GraphPad Prism 10.0. Biological replicates represent individual embryos, with sample sizes indicated in the figure legends. Each embryo was treated as one independent biological replicate. Control and CKO groups were compared using two-tailed unpaired Student’s t-tests, with P < 0.05 considered statistically significant. The initial morphometric analysis included six embryos per group. Power analysis based on palatal shelf cross-sectional area from this experiment indicated a minimum sample size of three embryos per group (80% power, two-sided α = 0.05), which was used as the minimum target for subsequent quantitative experiments.

## Results

### Neural crest-specific deletion of *Tmem2* impairs palatal shelf growth and causes cleft palate

To determine the role of TMEM2 in palatal development, we examined *Tmem2^f/f^; Sox10-Cre* embryos (CKO) and *Cre-negative* littermate controls. At E16.5, CKO embryos exhibited complete cleft palate, whereas no clefting was observed in controls (5/5 vs 0/5; Fig. 1A–E).

**Figure 1.**
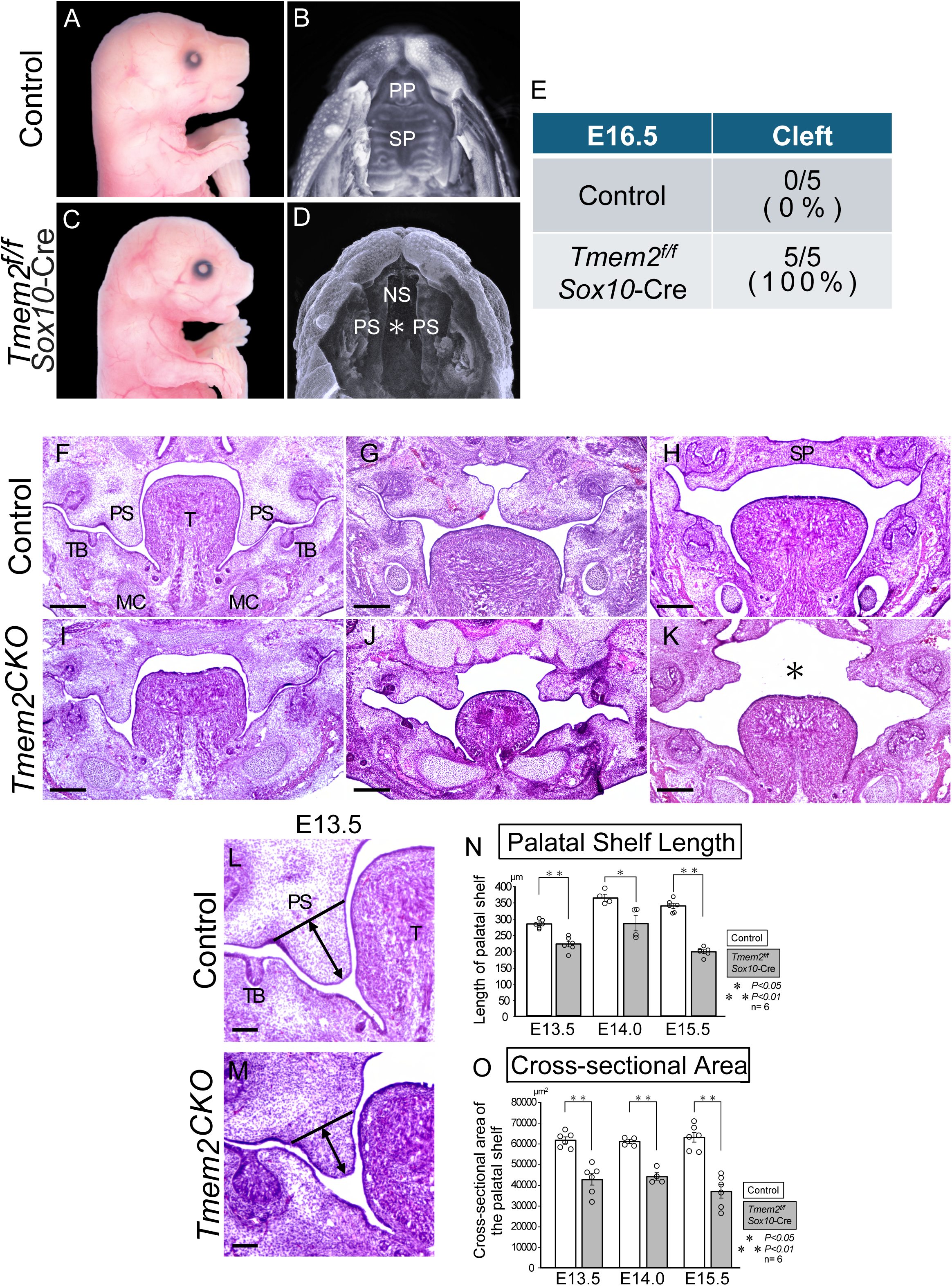
Neural crest-specific deletion of *Tmem2* impairs palatal shelf growth and causes cleft palate. (A–D) Gross morphology of *Cre-negative* litter mate control (A, B) and *Tmem2^f/f^; Sox10-Cre* (CKO) embryos (C, D) at E16.5. Lateral (A, C) and palatal views after mandibular removal (B, D) are shown. CKO embryos exhibited apparent shortening of the maxillary and mandibular regions, together with a complete cleft palate (asterisk in D). (E) Incidence of cleft palate at E16.5. All CKO embryos examined (5/5, 100%) exhibited cleft palate, whereas none of the controls did (0/5, 0%). (F–K) Representative hematoxylin and eosin (H&E)-stained coronal sections of palatal shelves from control (F–H) and CKO (I–K) embryos at E13.5 (F, I), E14.0 (G, J), and E15.5 (H, K). Palatal shelves were vertically oriented at E13.5 and elevated by E14.0 in both groups. CKO palatal shelves were consistently smaller than controls and remained unfused at E15.5 (asterisk in K). (L, M) Representative coronal sections at E13.5 illustrating the measurement of palatal shelf length and cross-sectional area in control (L) and CKO (M) embryos. Palatal shelf length was defined as the distance from the medial base to the lateral tip of the shelf (black double-headed arrow). (N, O) Quantitative analysis of palatal shelf length (N) and cross-sectional area (O) at E13.5, E14.0, and E15.5. Data are presented as mean ± SD (n = 6 embryos per group). P < 0.05, P < 0.01; Student’s t-test. Scale bars: 500 μm (F–K) and 50 μm (L, M). PP, primary palate; SP, secondary palate; NS, nasal septum; PS, palatal shelf; T, tongue; TB, tooth bud; MC, Meckel’s cartilage.

Histological analysis showed vertically oriented palatal shelves in both groups at E13.5 and elevated shelves by E14.0. However, CKO shelves were consistently smaller than controls and remained unfused at E15.5 (Fig. 1F–K). Quantitative analysis demonstrate significant reductions in palatal shelf length and cross-sectional area in CKO embryos at E13.5, E14.0, and E15.5 (Fig. 1N, O). Together, these findings indicate that neural crest-specific loss of Tmem2 impairs palatal shelf growth, while shelf elevation is preserved.

### CKO palatal shelves exhibit reduced proliferation and increased apoptosis

To examine the cellular changes accompanying reduced palatal shelf size, we assessed proliferation and apoptosis across the three developmental stages. Ki-67 immunostaining showed fewer proliferating cells in CKO palatal shelves at E14.0 (Fig. 2A, B), and quantitative analysis demonstrated a significant reduction in the proportion of Ki-67-positive cells in CKO palatal shelves at E13.5, E14.0, and E15.5 (Fig. 2C). Conversely, TUNEL staining at E14.0 revealed increased apoptotic cells in CKO palatal shelves (Fig. 2D, E), with significantly elevated TUNEL-positive cell numbers at all three developmental stages examined (Fig. 2F).

**Figure 2.**
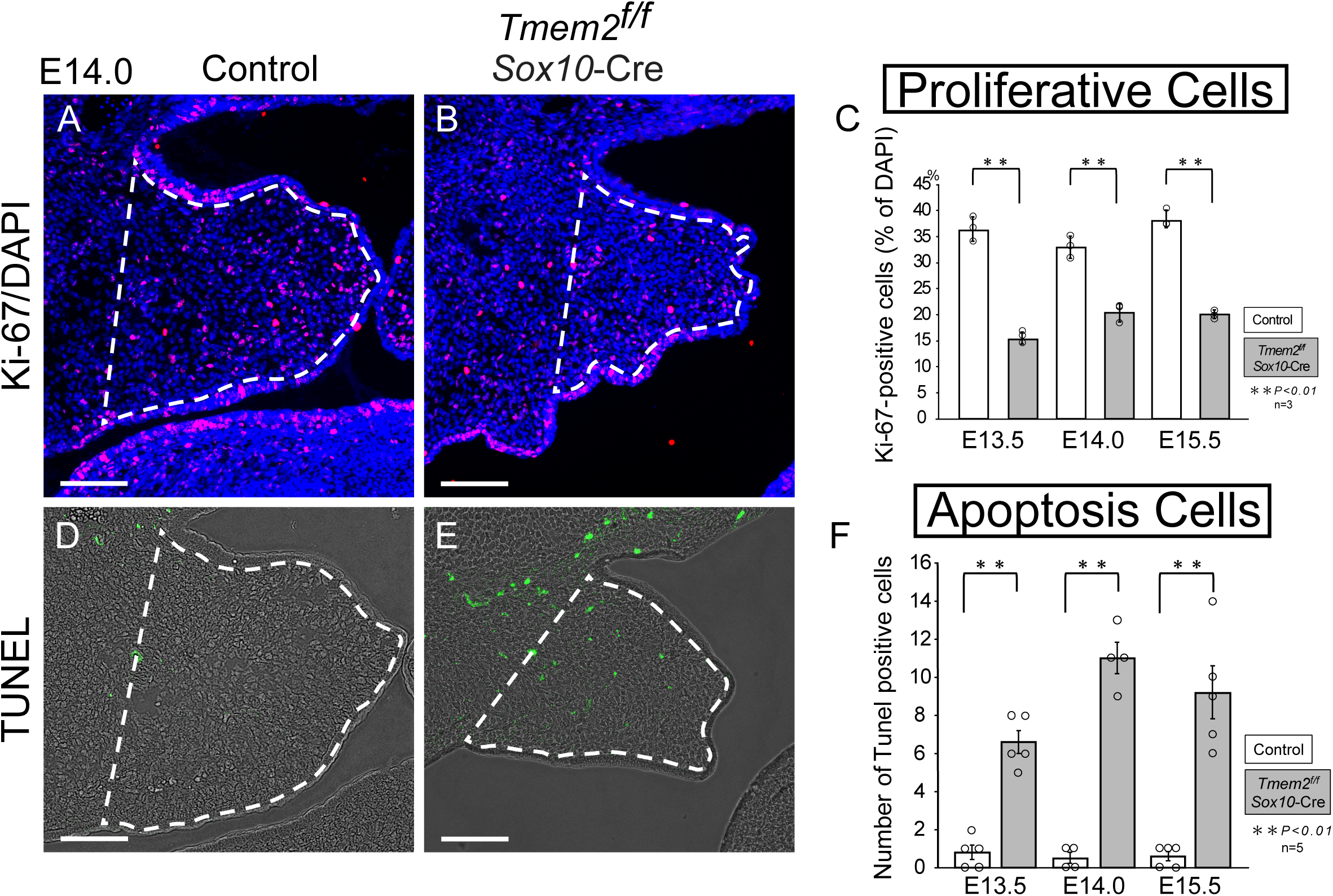
*Tmem2* deficiency is associated with reduced proliferation and increased apoptosis in the developing palate. (A, B) Representative Ki-67 immunofluorescence staining (red) with DAPI counterstaining (blue) in control (A) and CKO (B) palatal shelves at E14.0. (C) Quantification of the proportion of Ki-67-positive cells among total DAPI-positive cells at E13.5, E14.0, and E15.5. The proportion of Ki-67-positive cells was significantly reduced in CKO embryos at all developmental stages examined. (D, E) Representative TUNEL staining (green) with DAPI counterstaining (blue) in control (D) and CKO (E) palatal shelves at E14.0. (F) Quantification of TUNEL-positive cells at E13.5, E14.0, and E15.5. The number of TUNEL-positive cells was significantly increased in CKO embryos at all developmental stages examined. Data are presented as mean ± SD. Ki-67 analysis, n = 3 embryos per group; TUNEL analysis, n = 5, 4, and 5 embryos per group at E13.5, E14.0, and E15.5, respectively. P < 0.01; Student’s t-test. Scale bars: 100 μm.

### TMEM2 deficiency disrupts HA distribution and the spatial organization of NCC-derived cells in the palatal mesenchyme

To investigate the extracellular matrix changes accompanying impaired palatal shelf growth, we examined whether loss of the cell-surface hyaluronidase TMEM2 altered HA abundance and distribution in E13.5 palatal shelves. In control embryos, HABP staining revealed a spatially polarized pattern, with greater HA accumulation in the medial region and relatively lower levels in the lateral region (Fig. 3A). In contrast, CKO palatal shelves exhibited widespread HA accumulation throughout the mesenchyme (Fig. 3D). Quantitative analysis confirmed a significant increase in HABP-positive area and a reduction in the medial-to-lateral HA ratio in CKO embryos (Fig. 3G, H).

**Figure 3.**
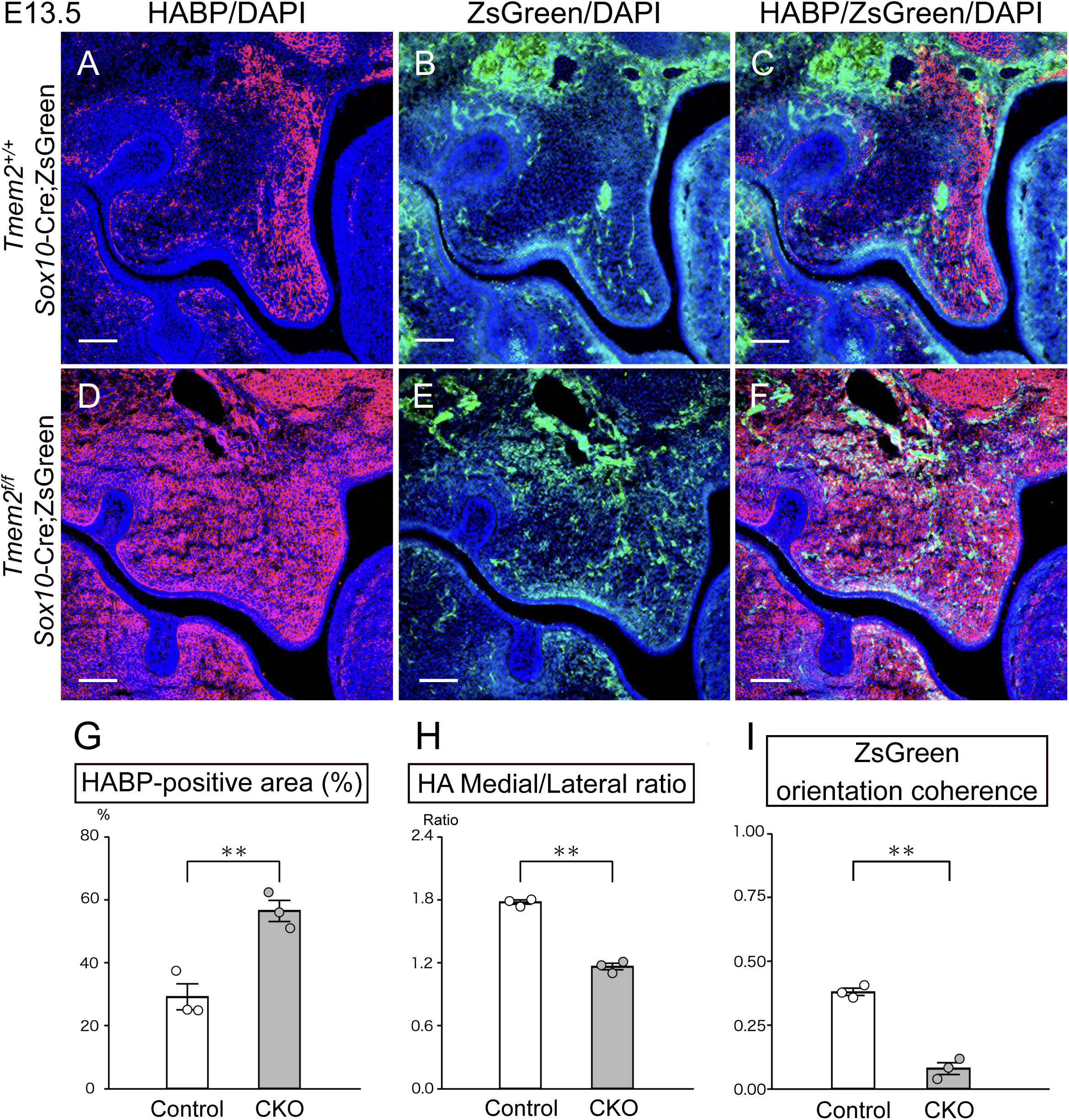
*Tmem2* deficiency disrupts HA distribution and the spatial organization of neural crest-derived cells in the developing palatal mesenchyme. (A–F) Representative coronal sections of E13.5 palatal shelves from control (*Tmem2^+/+^; Sox10-Cre*; ZsGreen) (A–C) and *Tmem2* CKO (*Tmem2^f/f^; Sox10-Cre*; ZsGreen) (D–F) embryos. HA was detected by HABP (red), NCC-derived cells were visualized by ZsGreen (green), and nuclei were counterstained with DAPI (blue). In control palatal shelves, HA showed a spatially polarized distribution, with greater accumulation in the medial region and relatively lower levels in the lateral region (A, C), while ZsGreen-positive NCC-derived cells exhibited a directionally organized pattern within the mesenchyme (B, C). In CKO embryos, HA accumulated extensively throughout the palatal mesenchyme, resulting in a more diffuse distribution (D, F), and ZsGreen-positive cells showed reduced directional organization (E, F). (G) Quantification of the HABP-positive area within the palatal mesenchyme. The HABP-positive area was significantly increased in CKO embryos compared with controls. (H) Quantification of the medial-to-lateral HA ratio. The medial-to-lateral ratio of HABP-positive area was significantly reduced in CKO embryos, indicating reduced spatial polarization of HA compared with controls. (I) Quantification of orientation coherence in ZsGreen-positive NCC-derived cells. Orientation coherence was significantly reduced in CKO embryos compared with controls. Data are presented as mean ± SD (n = 3 embryos per group). P < 0.01; Student’s t-test. Scale bars: 100 μm.

Previous work demonstrated a role for TMEM2-mediated HA remodeling in NCC migration (Inubushi et al. 2022). We therefore examined whether the altered HA distribution was accompanied by changes in the spatial organization of NCC-derived cells using ZsGreen lineage labeling. In control palatal shelves, ZsGreen-positive cells displayed a relatively organized and directionally coherent pattern within the mesenchyme (Fig. 3B, C). In CKO embryos, these cells exhibited reduced directional organization (Fig. 3E, F). Quantitative analysis confirmed significantly lower orientation coherence in CKO palatal shelves compared with controls (Fig. 3I). Together, these findings indicate that Tmem2 deficiency leads to widespread HA accumulation and reduced HA polarization and is associated with altered spatial organization of NCC-derived cells in the developing palatal mesenchyme.

### Transcriptomic analysis reveals enrichment of hypoxia-related pathways and suppression of cell-cycle programs in CKO palatal shelves

To identify molecular programs associated with the changes in HA distribution, NCC organization, and shelf growth, we profiled the transcriptomes of control and CKO palatal shelves at E14.0 by RNA sequencing. Differential expression analysis identified 2,027 DEGs, including 1,815 upregulated and 212 downregulated genes in CKO tissues (Fig. 4A).

**Figure 4.**
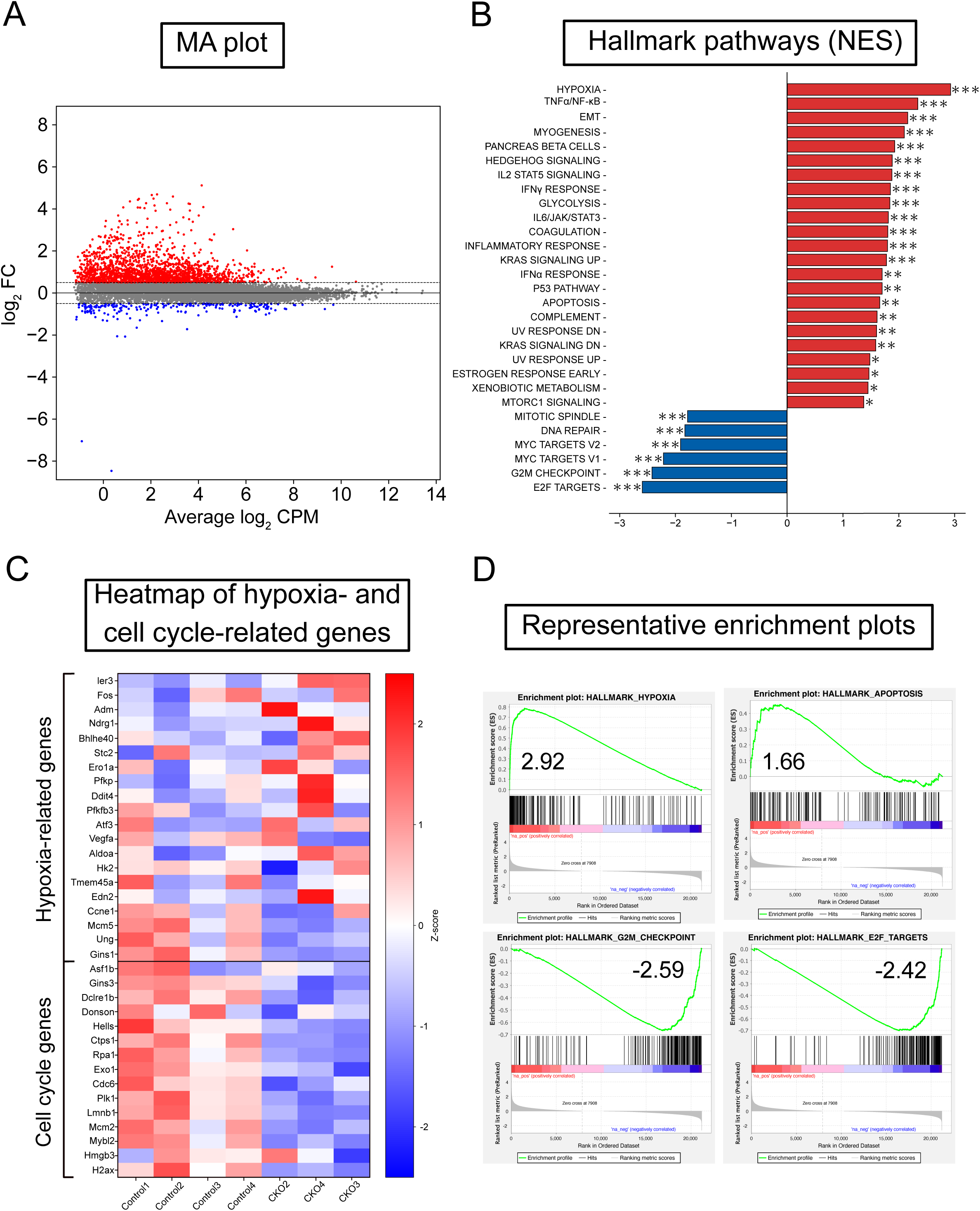
Transcriptomic analysis reveals enrichment of hypoxia-related responses and suppression of cell-cycle programs in Tmem2 CKO palatal tissues. (A) MA plot showing differential gene expression between control and CKO palatal tissues. Red and blue points indicate genes upregulated and downregulated, respectively, in CKO samples (|log2 fold change| > 0.5). (B) GSEA of Hallmark gene sets in control and CKO palatal tissues. Bars indicate normalized enrichment scores (NES). Positive and negative NES indicate enrichment in CKO and control palatal tissues, respectively. HALLMARK_HYPOXIA showed the highest positive NES among the significantly enriched gene sets, whereas cell-cycle-associated gene sets, including E2F targets and G2M checkpoint, were negatively enriched. Asterisks indicate significance based on FDR q-values: *q < 0.05, **q < 0.01, and ***q < 0.001. (C) Heatmap showing expression patterns of representative hypoxia-and cell cycle-related genes in control and CKO palatal tissues. Expression values are shown as Z-scores. (D) Representative GSEA enrichment plots showing positive enrichment of HALLMARK_HYPOXIA (NES = 2.92) and HALLMARK_APOPTOSIS (NES = 1.66), together with negative enrichment of HALLMARK_G2M_CHECKPOINT (NES = −2.59) and HALLMARK_E2F_TARGETS (NES = −2.42).

GSEA using Hallmark gene sets revealed marked enrichment of hypoxia-related signaling in CKO palatal tissues, with HALLMARK_HYPOXIA showing the highest positive normalized enrichment score (NES) among the significantly enriched pathways (Fig. 4B). Glycolysis and apoptosis were also positively enriched, whereas pathways associated with cell-cycle progression, including E2F targets and G2M checkpoint, were negatively enriched (Fig. 4B).

Consistent with these pathway-level changes, expression patterns of representative hypoxia- and cell cycle-related genes differed between control and CKO samples (Fig. 4C). Representative GSEA plots further demonstrated positive enrichment of hypoxia (NES = 2.92) and apoptosis (NES = 1.66), whereas G2M checkpoint (NES = −2.59) and E2F targets (NES = −2.42) were negatively enriched (Fig. 4D). Together, these transcriptomic findings indicate that *Tmem2* deficiency is associated with hypoxia- and apoptosis-related transcriptional responses and suppression of cell-cycle programs during palatal development.

### CKO palatal shelves show increased tissue hypoxia

Based on enrichment of hypoxia-related transcriptional signatures, we examined whether CKO palatal shelves exhibited evidence of tissue hypoxia at E13.5. HIF-1α-positive cells were significantly increased in CKO palatal shelves compared with controls (Fig. 5A–C). To further assess tissue hypoxia, pregnant mice were administered pimonidazole, which forms detectable adducts under hypoxic conditions. Pimonidazole-positive cells were also significantly increased in CKO palatal shelves (Fig. 5D–F). Together, these findings provide complementary evidence that loss of *Tmem2* in NCC-derived cells is associated with increased tissue hypoxia in the developing palate.

**Figure 5.**
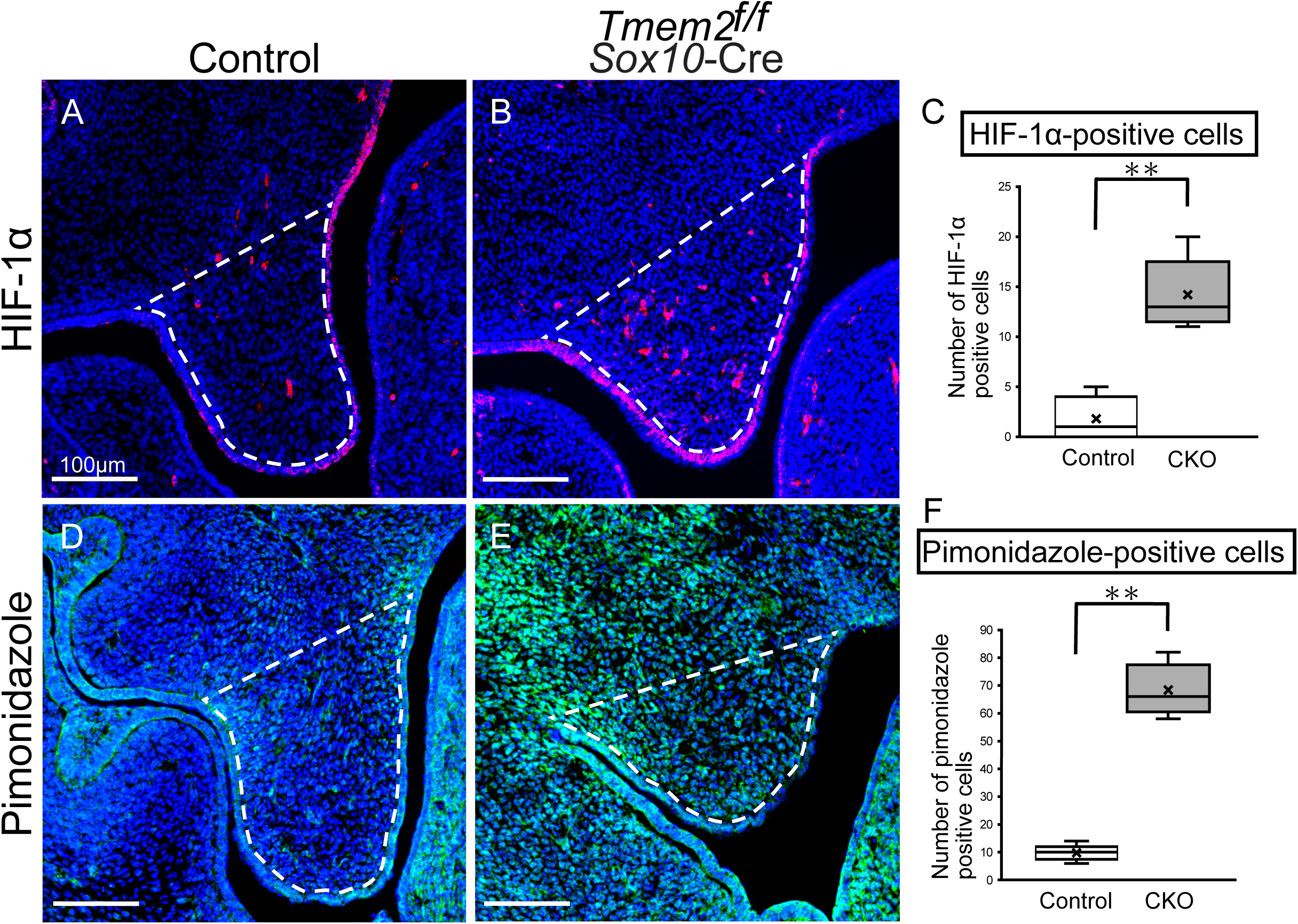
*Tmem2* deficiency is associated with increased tissue hypoxia in the developing palate. (A, B) Representative immunofluorescence staining for HIF-1α (red) and DAPI (blue) in E13.5 palatal shelves from control (A) and *Tmem2* CKO (B) embryos. Dashed lines outline the palatal mesenchymal region used for quantification. (C) Quantification of HIF-1α-positive cells within the palatal mesenchyme. The number of HIF-1α-positive cells was significantly increased in CKO embryos compared with controls. (D, E) Representative pimonidazole staining (red) with DAPI counterstaining (blue) in E13.5 palatal shelves from control (D) and CKO (E) embryos. Dashed lines outline the palatal mesenchymal region used for quantification. (F) Quantification of pimonidazole-positive cells within the palatal mesenchyme. The number of pimonidazole-positive cells was significantly increased in CKO embryos compared with controls. Data are presented as box-and-whisker plots. Boxes indicate the interquartile range, horizontal lines within the boxes indicate the median, whiskers indicate the minimum and maximum values, and crosses indicate the mean. HIF-1α and pimonidazole analyses, n = 5 embryos per group for each assay. **P < 0.01; Student’s t-test. Scale bars: 100 μm.

### *Tmem2* deficiency disrupts vascular and perivascular organization in the developing palate

To investigate whether tissue hypoxia coincided with abnormalities in the vessels that supply oxygen to the developing palate, we examined vascular organization at E13.5. Three-dimensional imaging of CD31-positive vessels at E13.5 showed an organized vascular network in control palatal shelves, with prominent interconnected vessels (Fig. 6A, B). In contrast, CKO palatal shelves exhibited visibly altered vascular architecture, with fewer prominent interconnected vessels and relatively finer vascular structures (Fig. 6C, D).

**Figure 6.**
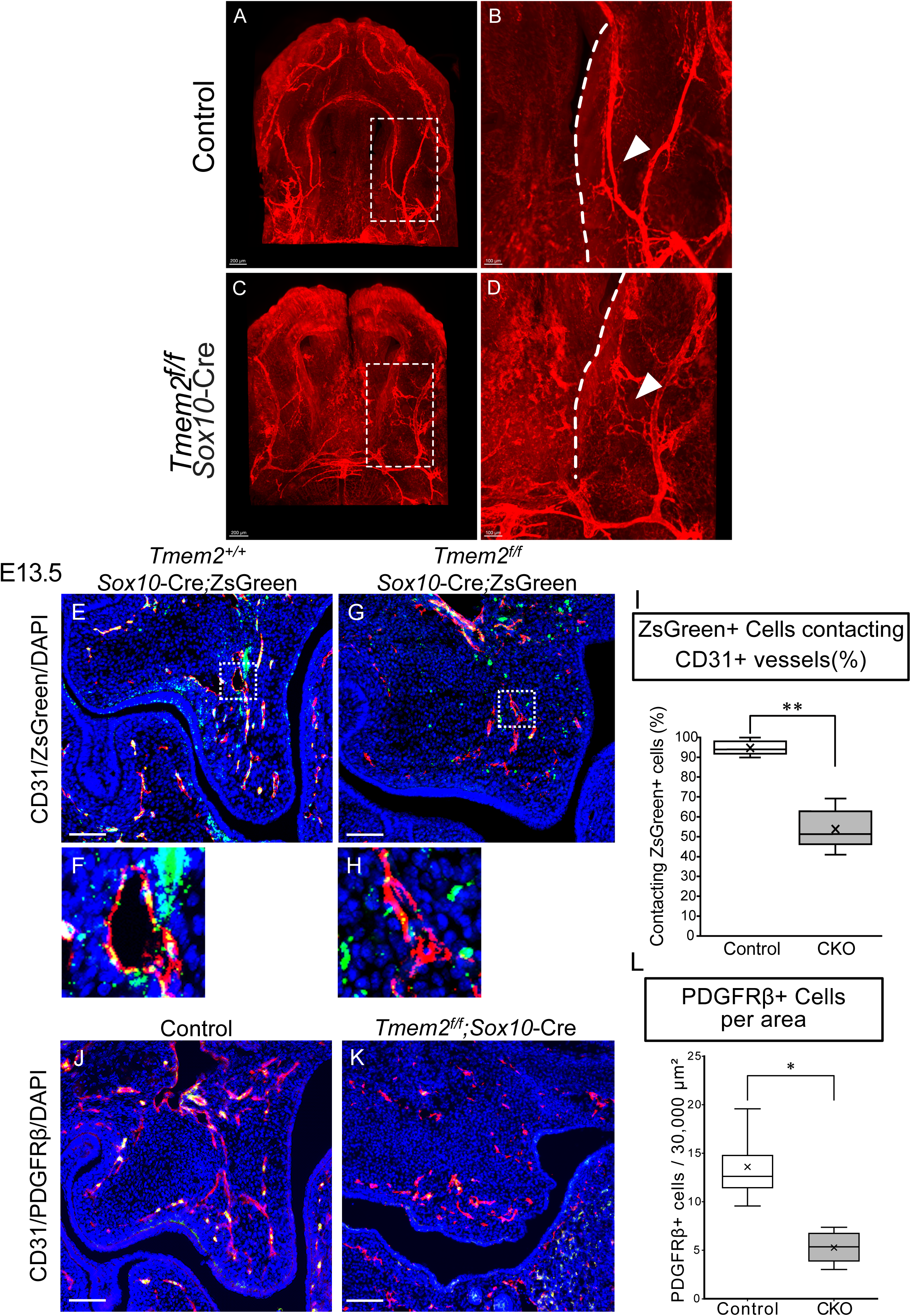
*Tmem2* deficiency disrupts vascular and perivascular organization in the developing palate. (A–D) Representative three-dimensional images of CD31-positive vasculature in E13.5 palatal tissues from control (A, B) and *Tmem2* CKO (C, D) embryos. Boxed regions in A and C are shown at higher magnification in B and D, respectively. Control palatal tissues exhibited prominent vessels with interconnected branches, whereas CKO tissues showed visibly altered vascular architecture, with fewer prominent interconnected vessels and relatively finer vascular structures. Arrowheads indicate representative vascular structures. Dashed lines indicate the approximate boundary of the palatal shelf. (E–H) Representative immunofluorescence images of CD31-positive vessels (red) and ZsGreen-positive NCC-derived cells (green) in E13.5 palatal shelves from control (E, F) and CKO (G, H) embryos. Nuclei were counterstained with DAPI (blue). Boxed regions in E and G are shown at higher magnification in F and H, respectively. (I) Quantification of the proportion of ZsGreen-positive cells contacting CD31-positive vessels. The proportion of ZsGreen-positive cells in contact with CD31-positive vessels was significantly reduced in CKO palatal shelves compared with controls. (J, K) Representative immunofluorescence staining for CD31 (red), PDGFRβ (green), and DAPI (blue) in E13.5 palatal shelves from control (J) and CKO (K) embryos. PDGFRβ-positive cells were less abundant, particularly in association with CD31-positive vessels, in CKO palatal shelves. (L) Quantification of PDGFRβ-positive cells per 30,000 μm² of palatal mesenchyme. The number of PDGFRβ-positive cells was significantly reduced in CKO embryos compared with controls. Data are presented as box-and-whisker plots. Boxes indicate the interquartile range, horizontal lines within the boxes indicate the median, whiskers indicate the minimum and maximum values, and crosses indicate the mean. For quantitative analyses, n = 5 embryos per group for CD31/ZsGreen analysis (I) and n = 4 embryos per group for PDGFRβ analysis (L). P < 0.05, P < 0.01; Student’s t-test. Scale bars: 200 μm (A, C), 100 μm (B, D), and 50 μm (E, G, J, K).

We next examined the spatial relationship between NCC-derived cells and the developing vasculature using *Sox10-Cre; ZsGreen* lineage labeling. In control palatal shelves, ZsGreen-positive NCC-derived cells were frequently observed in close association with CD31-positive vessels (Fig. 6E, F). This association was reduced in CKO embryos (Fig. 6G, H), and quantitative analysis confirmed a significant decrease in the proportion of ZsGreen-positive cells contacting CD31-positive vessels (Fig. 6I).

Because cranial NCCs can give rise to mural cells, including pericytes (Etchevers et al. 2001), we examined PDGFRβ-positive cells in the palatal mesenchyme. PDGFRβ-positive cells were frequently observed adjacent to CD31-positive vessels in controls, whereas fewer PDGFRβ-positive cells were detected in CKO palatal shelves (Fig. 6J, K). Quantification demonstrated a significant reduction in the number of PDGFRβ-positive cells per 30,000 μm² of palatal mesenchyme in CKO embryos (Fig. 6L). Together, these findings indicate that loss of *Tmem2* in NCC-derived cells is associated with altered vascular architecture, reduced vascular association of NCC-derived cells, and a decrease in PDGFRβ-positive mesenchymal cells in the developing palate.

## Discussion

Sox10-Cre-mediated *Tmem2* deletion allowed us to examine palatogenesis beyond the early embryonic lethality reported with Wnt1-Cre-mediated deletion (Inubushi et al. 2022). The CKO shelves elevated by E14.0 but remained smaller than control shelves and failed to fuse. Reduced proliferation and increased apoptosis were evident at each stage examined and may contribute to insufficient shelf growth. Together, these findings indicate that TMEM2-dependent regulation of the HA-rich extracellular environment contributes to the organization and growth of NCC-derived palatal mesenchyme.

Neural crest-specific deletion of *Has2* causes craniofacial abnormalities, including cleft palate (Lan et al. 2019), demonstrating the importance of HA synthesis in craniofacial development. Previous tissue-specific studies have further shown that *Tmem2* deficiency disrupts HA homeostasis in distinct craniofacial and skeletal tissues. Loss of *Tmem2* in K14-expressing epithelial cells impairs enamel formation (Nag et al. 2023), whereas chondrocyte-specific *Tmem2* deletion alters HA turnover and cartilage homeostasis (Murotani et al. 2026). Our findings extend these observations to palatal development by demonstrating a role for TMEM2-dependent HA degradation in NCC-derived mesenchyme. TMEM2 deficiency caused widespread HA accumulation and reduced medial-to-lateral polarization, suggesting that palatal morphogenesis depends not only on HA abundance but also on its spatial organization.

Altered HA distribution in CKO embryos was accompanied by reduced directional organization of NCC-derived cells. Previous work linked TMEM2 deficiency to impaired cranial NCC migration (Inubushi et al. 2022). TMEM2 has also been shown to promote integrin-dependent cell adhesion and migration by locally degrading HA at focal adhesion sites (Irie et al. 2021). Excess HA could alter cell–matrix interactions and thereby contribute to abnormal positioning and organization of NCC-derived cells.

The ZsGreen images capture spatial organization at a single stage, leaving cell migration and focal adhesion dynamics unresolved.

Transcriptome profiling identified hypoxia as the Hallmark gene set with the highest positive enrichment score among the significant results. The glycolysis signature is consistent with the metabolic response to reduced oxygen availability (Kim et al. 2006; Semenza 2012). Increased HIF-1α and pimonidazole labeling provided complementary evidence of tissue hypoxia. Experimental studies link embryonic hypoxia or hypoxia-related injury with susceptibility to orofacial clefting, including cleft palate (Azarbayjani and Danielsson 2001; Millicovsky and Johnston 1981). These observations motivated examination of the palatal vasculature as a possible contributor to the hypoxic phenotype.

Vascular remodeling accompanies normal palatal shelf elevation and fusion (Amin et al. 1994), and a relationship between TMEM2-dependent HA turnover and developmental angiogenesis has also been demonstrated in zebrafish, where loss of *Tmem2* causes excessive HA accumulation around developing vessels, altered Vegf signaling, and impaired angiogenesis (De Angelis et al. 2017). Our findings extend this relationship to mammalian palatal development, suggesting that TMEM2-dependent HA remodeling in NCC-derived mesenchyme may support proper vascular and perivascular organization.

Cranial NCCs contribute to pericytes and vascular smooth muscle cells of craniofacial vessels (Etchevers et al. 2001), and perivascular cells are essential for vascular morphogenesis and stabilization (Armulik et al. 2011; Hellström et al. 2001). NCC-derived VEGFA supports vascular development in craniofacial tissues, with studies in the developing jaw providing evidence for a paracrine mechanism (Wiszniak et al. 2015). In the developing palate, Wnt1-Cre-mediated deletion of Vegfa reduces palatal shelf vascularization and is accompanied by reduced cell proliferation, abnormal shelf elongation and elevation, and cleft palate (Hill et al. 2015). Together, these studies support the concept that NCC-derived cells regulate craniofacial vascular development through both perivascular and paracrine mechanisms.

Consistent with this relationship, NCC-derived cells were frequently associated with CD31-positive vessels in control palatal shelves, whereas this association was significantly reduced in CKO embryos. CKO palatal shelves also contained fewer PDGFRβ-positive cells per unit mesenchymal area and exhibited altered vascular architecture. Because *Sox10-Cre* targets the NCC lineage rather than vascular endothelial cells, these abnormalities are unlikely to result primarily from direct loss of TMEM2 in endothelial cells. Instead, abnormal HA accumulation and disruption of NCC organization may compromise the formation and maintenance of the perivascular microenvironment. One possibility is that excessive HA impairs the migration or positioning of NCC-derived cells toward developing vessels. Alternatively, TMEM2 deficiency may interfere with the acquisition or maintenance of a perivascular/mural-cell phenotype after NCC-derived cells reach the vascular niche.

The present data do not distinguish whether the reduction in PDGFRβ-positive cells primarily reflects defective perivascular positioning, altered acquisition or maintenance of a mural-cell phenotype, or both.

The coexistence of vascular abnormalities and tissue hypoxia raises the possibility that oxygen delivery is impaired in CKO palatal shelves. The CD31 images describe vessel architecture, and whether perfusion or oxygen delivery was reduced remains open. Hypoxia could contribute to the proliferation and survival defects, alongside changes in the HA-rich matrix and mesenchymal organization. The relative contributions of these changes to insufficient shelf growth remain to be determined.

Three-dimensional CD31 imaging revealed qualitative differences in vascular architecture. Quantifying vessel diameter, branching, and connectivity would define those changes more precisely. Lineage tracing combined with complementary mural-cell markers would help establish the identity and fate of the affected cells and distinguish altered perivascular positioning from impaired mural-cell differentiation or maintenance. Functional studies would be needed to establish whether altered oxygen delivery contributes to the growth phenotype.

In conclusion, loss of Tmem2 in the neural crest lineage causes cleft palate with impaired palatal shelf growth despite preserved elevation. TMEM2 deficiency disrupts HA distribution and NCC-derived mesenchymal organization and is accompanied by vascular and perivascular abnormalities and tissue hypoxia. These findings support a role for TMEM2-dependent HA remodeling in coordinating the extracellular and cellular organization required for normal palatal development.

## Author Contributions

H. Shamma contributed to data acquisition, analysis, and interpretation, and drafted the manuscript; T. Inubushi contributed to conception, design, data acquisition, analysis, and interpretation, and critically revised the manuscript; J.I. Sasaki, H. Sakai, T. Murotani, Y. Nakanishi, and T. Tomohiro contributed to analysis and interpretation, and critically revised the manuscript; H. Kurosaka, Y. Yamaguchi and T. Yamashiro contributed to interpretation and critically revised the manuscript. All authors gave final approval and agree to be accountable for all aspects of the work.

## Acknowledgments

We thank Ms. Mayumi Yoshimoto for the excellent care and maintenance of our mouse colony. This work was funded by Grants-in-Aid for Scientific Research (24K02652 to T. Inubushi) from the Japan Society for the Promotion of Science and by the Japan Science and Technology Agency FOREST Program (JPMJFR220J to T.I.).

## Data availability

All relevant data can be found within the article and its supplementary information.

